# Exploiting the mediating role of the metabolome to unravel transcript-to-phenotype associations

**DOI:** 10.1101/2022.06.08.495285

**Authors:** Chiara Auwerx, Marie C. Sadler, Alexandre Reymond, Zoltán Kutalik, Eleonora Porcu

**Affiliations:** Center for Integrative Genomics, University of Lausanne, Lausanne 1015, Switzerland; Swiss Institute of Bioinformatics, Lausanne 1015, Switzerland; University Center for Primary Care and Public Health, Lausanne 1010, Switzerland; Department of Computational Biology, University of Lausanne, Lausanne 1015, Switzerland

## Abstract

Despite the success of genome-wide association studies (GWASs) in identifying genetic variants associated with complex traits, understanding the mechanisms behind these statistical associations remains challenging. Several methods that integrate methylation, gene expression, and protein quantitative trait loci (QTLs) with GWAS data to determine their causal role in the path from genotype to phenotype have been proposed. Here, we developed and applied a multi-omics Mendelian randomization (MR) framework to study how metabolites mediate the effect of gene expression on complex traits. We identified 206 transcript-metabolite-trait causal triplets for 28 medically relevant phenotypes. Sixty-seven of these associations were missed by classical transcriptome-wide MR, which only uses gene expression and GWAS data. Among these, we identify biologically relevant pathways, such as between *ANKH* and calcium levels mediated by citrate and *SLC6A12* and serum creatinine through modulation of the levels of the renal osmolyte betaine. We show that the signals missed by transcriptome-wide MR are found thanks to the gain in power allowed by integrating multiple omics-layer. Simulation analyses show that with larger molecular QTL studies and in case of mediated effects, our multi-omics MR framework outperforms classical MR approaches designed to detect causal relationships between single molecular traits and complex phenotypes.

## Introduction

Genome-wide association studies (GWAS) have identified thousands of single nucleotide polymorphisms (SNPs) associated with a wide range of complex traits [1, 2]. However, the path from GWAS to biology is not straightforward as most SNPs implicated by GWASs reside in non-coding regions of the genome [1] and do not directly inform on the functional mechanism through which variants exert their effect on phenotypes.

GWASs have been performed on gene expression [3], DNA methylation [4], protein [5], and metabolites [6, 7] levels, identifying genetic variants influencing molecular traits, commonly referred to as molecular quantitative trait loci (molQTLs). The large overlap between complex and molecular trait-associated variants suggests that integrating these data can help interpreting GWAS loci [8-10]. Advances in the field of transcriptomics make gene expression the best studied molecular phenotype, thanks to the presence of large expression QTL (eQTL) studies (e.g., eQTLGen Consortium [3], N > 30,000). Availability of these datasets fostered the development of summary statistic-based statistical approaches aiming at identifying associations between transcripts and complex traits [11-14], prioritizing genes from known GWAS loci for functional follow-up, and inferring the directionality of these relations [12, 15]. However, the cascade of events that mediates the effect of genetic variants on complex traits involves more than one molecular trait. Although approaches used for gene expression can be extended to other molecular data, investigating whether these molecular traits reside along the same causal pathway remains under-explored and only recently studies applied colocalization and Mendelian randomization (MR) to methylation, gene expression, and protein levels data [16-19] and to a lesser extent to metabolic QTLs (mQTL).

Metabolites are often the final products of cellular regulatory processes and the most proximal omic layer to complex phenotypes. Their levels could thus represent the ultimate response of biological systems to genetic and environmental changes. For instance, the metabolic status of organisms reflects disease progression, as metabolic disturbances can often be observed several years prior to the symptomatic phase [20-22]. Therefore, using metabolomics to identify early-stage biomarkers of complex phenotypes, such as prediabetes and COVID-19 susceptibility, has gained increased interest [23, 24]. While two-sample MR approaches using metabolites as single exposure have revealed biomarkers for several diseases [25-27], these analyses focused on the prediction of disease risk rather than on deciphering the mechanisms of discovered associations.

In an MR framework, when hypothesizing a mediating role for the metabolome on the genotype-to-phenotype axis, the primary exposure may be defined as an upstream omic layer, such as for instance gene expression. Integrating transcriptomics with metabolomics data can provide insights into how metabolites are regulated, elucidating targetable functional mechanisms. To explore this scenario, we developed an integrative MR analysis combining summary-level multi-omics data to compute the indirect effect of gene expression on complex traits mediated by metabolites. Our integrative analysis of GWAS, eQTL, and mQTL data consists of three steps (Figure 1). First, we map the transcriptome to the metabolome by identifying causal associations between transcripts and metabolites. Next, we screen the metabolites for downstream causal effects on 28 complex phenotypes, resulting in the identification of gene expression → metabolite → phenotype cascades. (Figure 1A). In parallel, we prioritize trait-associated genes by testing the association of transcripts with phenotypes (Figure 1B). Third, for transcripts identified in either (a) or (b) we test whether the identified target genes exert their effect on the phenotype through the metabolite using multivariable MR (MVMR; Figure 1C). Finally, we carried out extensive power analyses to determine under which conditions the mediation analysis (Figure 1C) outperforms the conventional exposure-outcome MR framework (Figure 1B).

**Figure 1.**
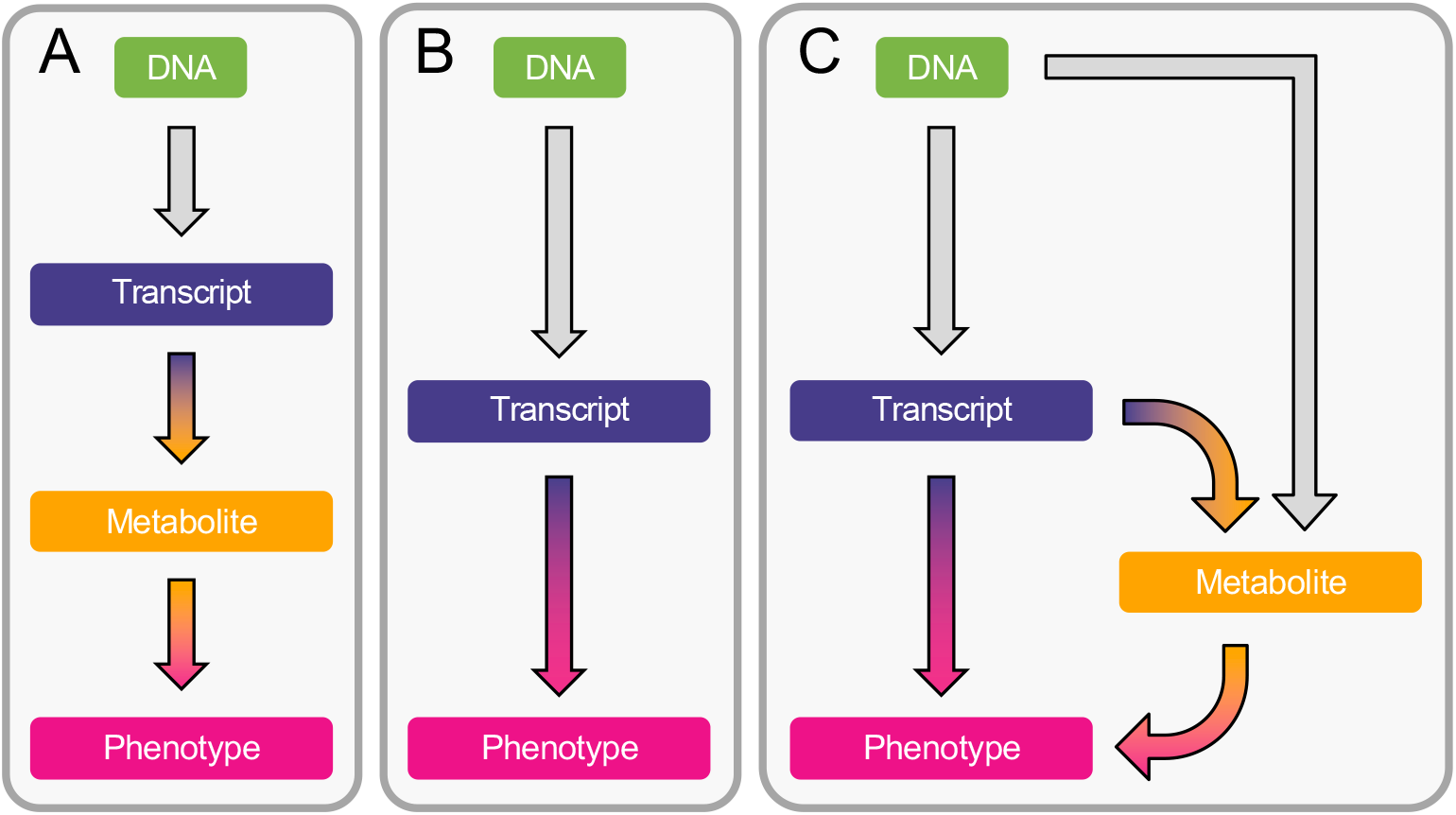
Methods overview. **A**. Estimation of the causal effect from transcript to metabolite and from metabolite to phenotype through univariable Mendelian randomization (MR). **B**. Estimation of the causal effect from transcript to phenotype through univariable transcriptome-wide MR (TWMR). **C**. Estimation of the direct (i.e., not mediated by the metabolites) and mediated effect of transcripts on phenotypes through multivariable MR (MVMR) by accounting for mediation through the metabolome.

## Results

### Mapping the transcriptome onto the metabolome

We applied univariable MR to identify metabolites whose levels are causally influenced by transcript levels in whole blood (Figure 1A). Summary statistics for *cis*-eQTLs stem from the eQTLGen Consortium metanalysis of 19,942 transcripts in 31,684 individuals [3], while summary statistics for mQTLs originate from a metanalysis of 453 metabolites in 7,824 individuals from two independent European cohorts: TwinsUK (N = 6,056) and KORA (N = 1,768) [6]. After selecting SNPs included in both datasets, our analysis was restricted to 7,883 transcripts with ≥ 3 instrumental variables (IVs) (see Methods). By testing each gene for association with the 453 metabolites, we detected 191 genes whose transcript levels causally impacted 154 metabolites, resulting in 257 unique transcript-metabolite associations (*P* < 0.05⁄7,883 = 6.3 × 10^−06^ ; Supplemental Table 1). Overall, 83% of the involved genes (159/191) were causally influencing the level of a single metabolite, while *TMEM258* and *FADS2* affected 12 metabolites.

### Mapping the metabolome onto complex phenotypes

Univariable metabolome-wide MR (MWMR) was used to identify causal relationships between 87 metabolites with ≥ 3 IVs and 28 complex phenotypes, including anthropometric traits, cardiovascular assessments, and blood biomarkers (Figure 1A, Supplemental Table 2). Phenotype summary statistics originate from the UK biobank (UKB) [28]. Overall, 54 metabolites were associated with at least one phenotype (*P* < 0.05⁄87 = 5.7 × 10^−04^), resulting in 133 unique metabolite-phenotype associations (Supplemental Table 3).

### Mapping the transcriptome onto complex phenotypes

We applied univariable transcriptome-wide MR (TWMR) to identify associations between expression levels of 10,435 transcripts from the eQTLGen Consortium with ≥ 3 IVs measured in both exposure and outcome datasets and the same 28 UKB phenotypes described in the previous section (Figure 1B). In total, 1,659 transcripts associated with at least one phenotype (*P* < 0.05⁄10,435 = 4.8 × 10^−06^), resulting in 3,168 unique transcript-phenotype associations (Supplemental Table 4).

### Mapping metabolome-mediated effects of the transcriptome onto complex phenotypes

The mapping of putative causal transcripts and metabolites performed in the previous steps provides the opportunity to infer the mediating role of the metabolome in biological processes leading to transcript-phenotype associations. We combined the 257 transcript-metabolite and 133 metabolite-trait significant associations to pinpoint 206 transcript-metabolite-phenotype causal triplets (Supplemental Table 5). For each of these putative mechanisms, we applied a multivariable MR (MVMR) approach to compute the direct effect of gene expression on the phenotype (see Methods; Figure 1C). Regressing the total effect (*α*_*TP*_) on the direct effect (*α*_*d*_) (Figure 2A), we estimated that for our 206 mediated associations, 79% [95% CI: 72%-86%] of the transcript effect on the phenotype was direct and thus not mediated by the metabolites (Figure 2B).

**Figure 2.**
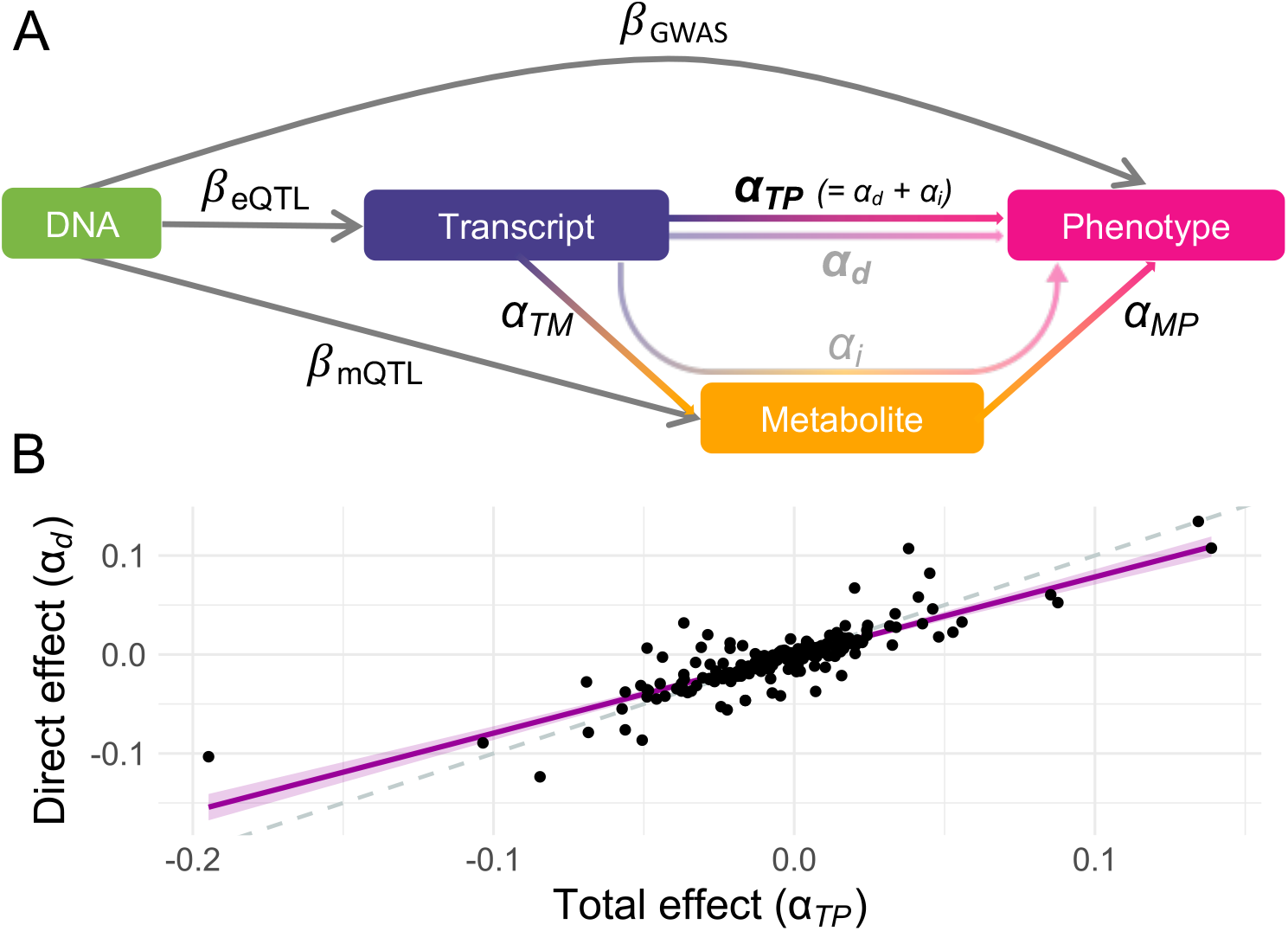
Direct and mediated effects. **A**. Graphical representation of the multivariable Mendelian randomization (MVMR) framework for mediation analysis: DNA represents genetic instrumental variables (IVs) chosen to be directly associated with either the exposure (transcript; *β*_*eQTL*_) or the mediator (metabolite; *β*_*mQTL*_) through summary statistics. The effect of these IVs on the outcome (phenotype; *β*_*GWAS*_) originate from genome-wide association studies (GWASs) summary statistics. Total effects *α*_*TP*_ of transcripts on phenotypes are estimated by transcriptome-wide Mendelian randomization (TWMR), while direct effects *α*_*d*_ are estimated by MVMR. Total effects *α*_*TP*_ are assumed to equal the sum of the direct *α*_*d*_ and indirect *α*_*i*_ (i.e., mediated) effects, the two former being shown in bold and depicted in **B. B**. Direct (*α*_*d*_; y-axis) and total (*α*_*TP*_; x-axis) effects for the 206 transcript-metabolite-trait causal triplets. The dashed line represents the identity, while the pink line represents the regression line with a shaded 95% confidence interval.

### Molecular mechanisms of genotype-to-phenotype associations

Dissecting causal triplets allows gaining mechanistic insights into biological pathways linking genes to phenotypes. For instance, expression of *TMEM258* [MIM: 617615], *FADS1* [MIM: 606148], and *FADS2* [MIM: 606149], all mapping to a region on chromosome 11 (Figure 3A), were found to influence a total of 12 complex phenotypes through modulation of 1-arachidonoylglycerophosphocholine (LPC(20:4); HMDB0010395; *α*_*TMEM258*_ = −1.02 ; *P* = 8.0 × 10^−81^ ; *α*_*FADS*1_ = −0.39; *P* = 4.6 × 10^−15^ ; *α*_*FADS*2_ = −0.63; *P* = 5.1 × 10^−62^) and 1-arachidonoylglycerophosphoethanolamine (LPE(20:4); HMDB0011517; *α*_*TMEM*258_ = −0.68; *P* = 1.1 × 10^−37^ ; *α*_*FADS*1_ = −0.30; *P* = 1.4 × 10^−07^ ; *α*_*FADS*2_ = −0.37; *P* = 1.2 × 10^−18^) levels (Figure 3B-C the known pleiotropy of the region (i.e., > 6,000 associations reported in the GWAS Catalog as of May 2022). Interestingly, involved metabolites are complex lipids synthesized from arachidonic acid, a product of the rate-limiting enzymes encoded by *FADS1* and *FADS2* (Figure 3B). Recently, polymorphisms affecting the expression of these genes were shown to associate with the levels of over 50 complex lipids, including the ones identified by our study [29]. Overall, this example illustrates how our method can capture meaningful biological associations and shed light on underlying molecular pathways of pleiotropy.

**Figure 3.**
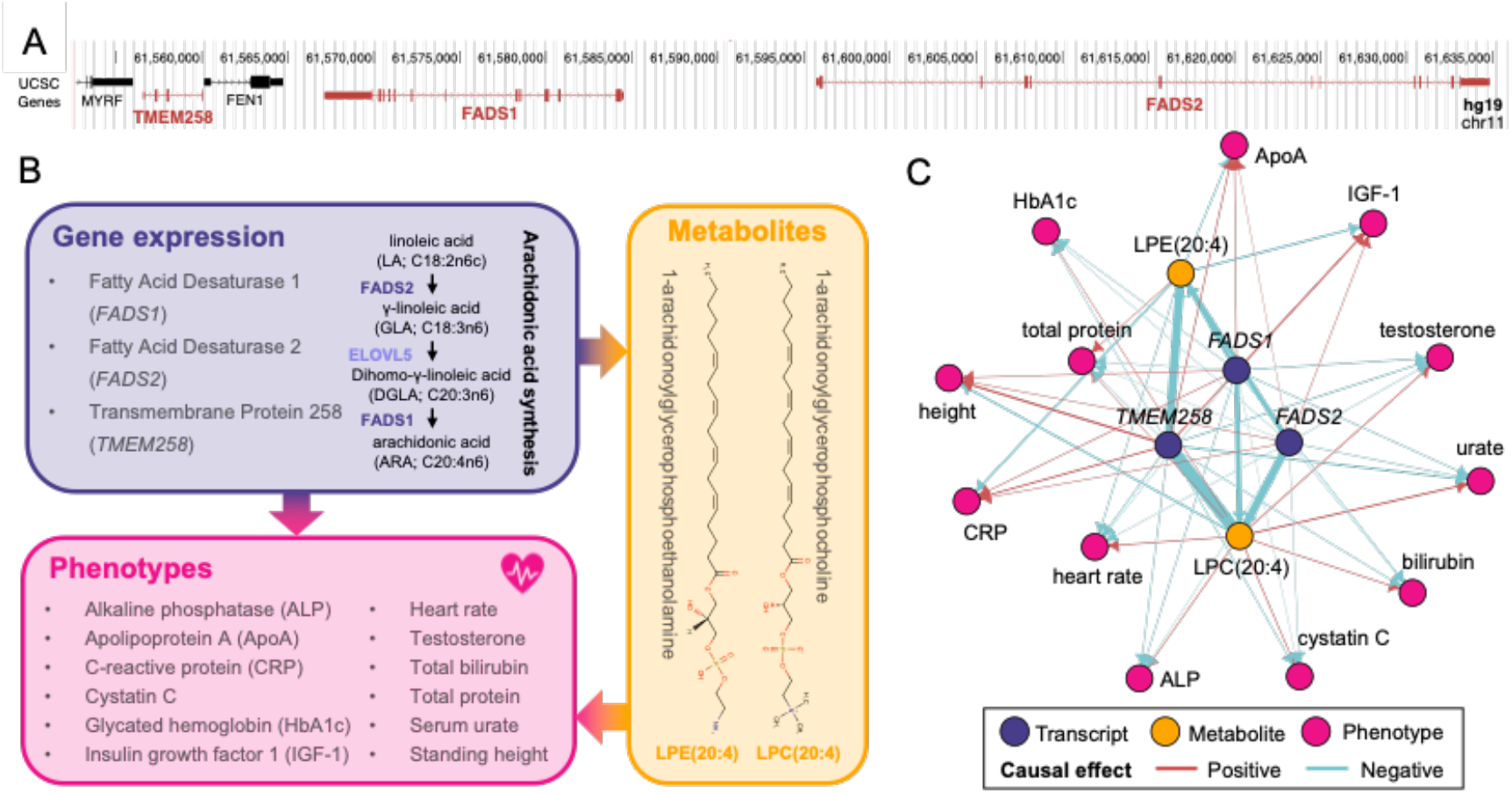
**A**. Genome browser (hg19) view of the genomic region on chromosome 11 encompassing *TMEM258, FADS1*, and *FADS2* (red). **B**. Diagram of the mediation signals detected for *TMEM258, FADS1*, and *FADS2*. Two of the implicated genes encode enzymes involved in arachidonic synthesis (purple). Involved genes impact 12 phenotypes (pink) through alteration of the levels of two metabolites, 1-arachidonoylglycerophosphocholine (LPC(20:4)) and 1-arachidonoylglycerophosphoethanolamine (LPE(20:4)), whose structure is depicted (orange). **C**. Network of the 42 transcript-metabolite-trait causal triplets involving *TMEM258, FADS1*, and *FADS2*. Nodes represent genes (purple), metabolites (orange), or phenotypes (pink). Edges indicate the direction of the effects estimated through univariable Mendelian randomization. Width of edges is proportional to effect size and color indicates if the effect is positive (red) or negative (blue).

### Power analysis

Importantly, only 33% (67/206) of the causal triplets showed a significant total transcript-to-phenotype effect (i.e., estimated by TWMR), suggesting that the method lacks power under current settings. To characterize the parameter regime where the power to detect indirect effects is larger than it is for total effects, we performed simulations using different settings for the mediated effect. We simulated 1,000 scenarios where a transcript with 6% heritability (i.e., median *h*^2^ in the eQTLGen data) has a causal effect of 0.035 (i.e., ∼65% of power in TWMR at α = 0.05) on a phenotype (see Methods). We varied two parameters characterizing the mediation:

a. the proportion (*ρ*) of direct (*α*_*d*_) to total (*α*_*TP*_) effect (i.e., effect not mediated by the metabolite) from 0 to 1;
b. the ratio (*σ*) between the transcript-to-metabolite (*α*_*TM*_) and the metabolite-to-phenotype (*α*_*MP*_) effects, exploring the range from 0.1 to 10.

Simulations show that with current sample sizes (i.e., *N*_*GWAS*_ = 300,000, *N*_*eQTL*_ = 32,000, and *N*_*mQTL*_= 8,000), when *α*_*MP*_ > *α*_*TM*_ (i.e., *σ* < 1), TWMR has increased power to detect significant transcript-to-phenotype associations over the full range of proportion of mediated effect *ρ* (Figure 4A, Supplemental Table 6). However, for all 206 causal triplets, we observed *σ* > 1 (Supplemental Figure 1). Under this condition, and assuming that the total effect of the transcript on the phenotype is dominated by the effect mediated by the metabolite (i.e., *ρ* < 0.5), TWMR had less power than the approach identifying mediators (Figure 4A, Supplemental Table 6), confirming that significant associations were missed by TWMR due to power issues related to the proportion of mediated effect.

**Figure 4.**
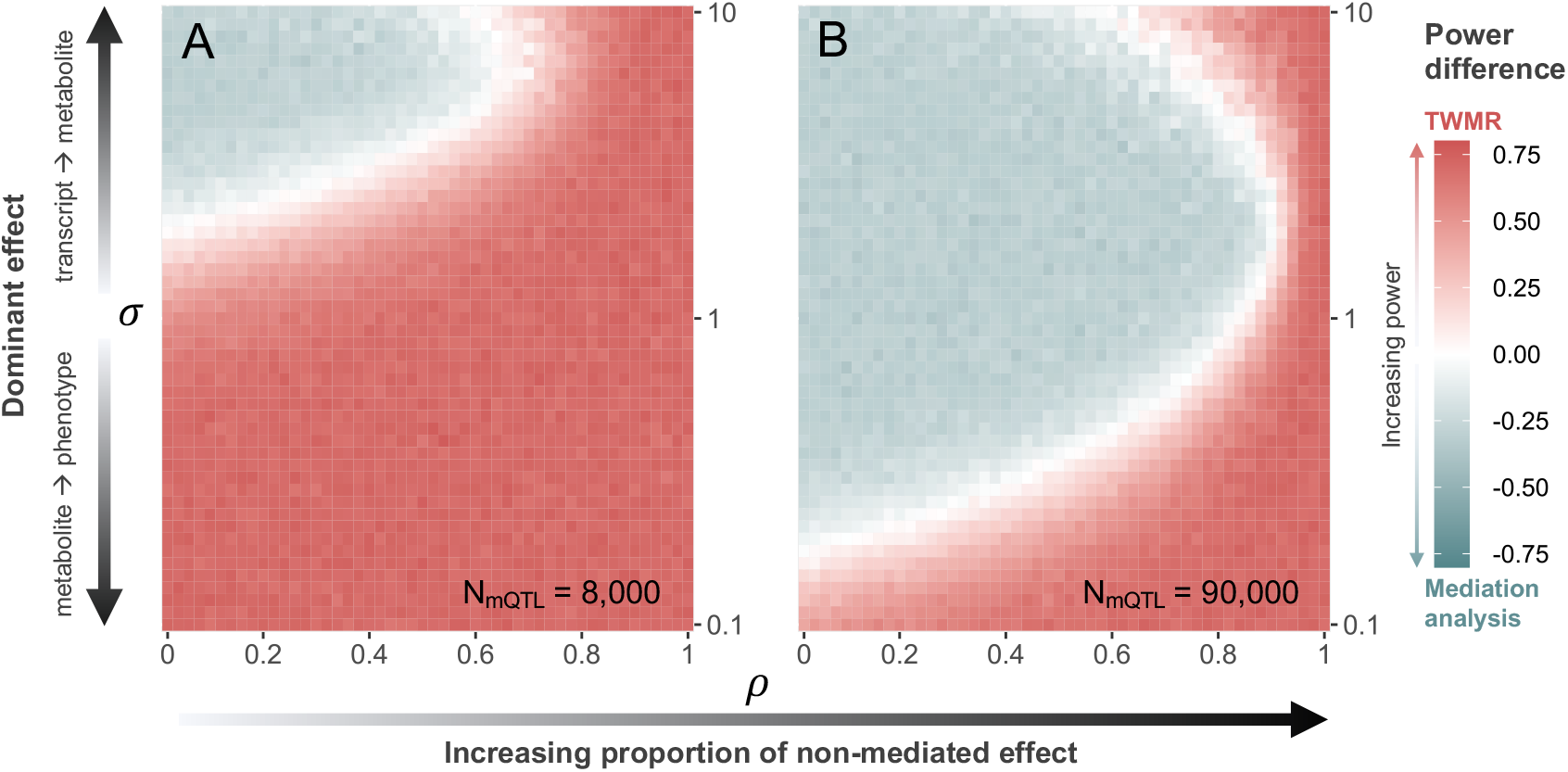
Heatmap showing the difference in statistical power between transcriptome-wide Mendelian randomization (TWMR) and mediation analysis through multivariable Mendelian randomization (MVMR) at current (**A**; N = 8,000) and realistic future (**B**; N = 90 000) mQTL dataset sample sizes. The x-axis shows the proportion (*ρ*) of direct (*α*_*d*_) to total (*α*_*TP*_) effect (i.e., effect not mediated by the metabolite) ranging from 0 to 1. The y-axis shows the ratio (*σ*) between the transcript-to-metabolite (*α*_*TM*_) and the metabolite-to-phenotype (*α*_*MP*_) effects, ranging from 0.1 to 10. Red vs. blue indicates higher power for TWMR vs. mediation analysis, respectively, while white represents equal power between the two approaches.

Repeating the simulations with a mQTL sample size of 90,000, nearing state-of-the-art sample sizes [7], we observe a strong shift in the above-described trends (Figure 4B, Supplemental Table 7). Specifically, when the effect of the transcript on the phenotype is dominated by the effect mediated by the metabolite (*ρ* < 0.3), mediation analysis has more power than TWMR when *σ* > 0.2. For larger proportions of direct effect, TWMR has increased power the more *σ* differs from 1.

### Identifying new genotype-to-phenotype associations

The 139 triplets that were not identified through TWMR due to power issues represent putative new causal relations. For instance, we observed that *ANKH* [MIM: 605145] expression decreased citrate levels (*α*_*ANKH*_ = −0.30; *P* = 2.2 × 10^−06^), which itself increased serum calcium levels (*α*_*citrate*_ = 0.07; *P* = 6.5 × 10^−10^), despite the lack of a significant TWMR effect of *ANKH* expression on calcium levels (*α*_*ANKH*_ = −0.02; *P* = 0.03). Citrate has a high binding affinity for calcium and influences its bioavailability by complexing calcium-phosphate during extracellular matrix mineralization and releasing calcium during bone resorption [30]. *ANKH* encodes for a transmembrane protein that channels inorganic pyrophosphate to the extracellular matrix where at low concentrations, it inhibits mineralization [31]. Accordingly, mutations in the gene have been associated with several rare mineralization disorders [MIM: 123000, 118600] [32]. Together, our data support the role of *ANKH* in calcium homeostasis through regulation of citrate levels.

In another example, *SLC6A12* [MIM: 603080], which encodes the Betaine/GABA Transporter-1 (*BGT-1*) involved in betaine and GABA uptake [33], was identified as a negative regulator of betaine (*α*_*SLC*6*A*12_ = −0.37 ; *P* = 8.2 × 10^−08^). While blood betaine levels negatively impacted serum creatinine levels (*α*_*betaine*_ = −0.06; *P* = 1.7 × 10^−07^), the effect of *SLC6A12* expression on creatinine was not significant (*α*_*SLC*6*A*12_ = 0.02; *P* = 1.5 × 10^−03^). This observation is particularly interesting given that betaine acts as a protective renal osmolyte whose plasma and kidney tissue concentration were found to be downregulated in renal ischemia/reperfusion injury [34, 35] and whose urine levels have been proposed as a biomarker for chronic kidney disease progression [36]. As both renal conditions are commonly monitored through serum creatinine levels, our data support the critical role of osmolyte homeostasis in renal health.

## Discussion

In this study, we combined MR approaches integrating eQTL, mQTL, and GWAS summary statistics to explore the role of the metabolome in mediating the effect of the transcriptome on complex phenotypes. Applied to 28 medically relevant traits, our approach revealed 206 causal transcript-metabolite-phenotype triplets. Among the 67 signals that were also identified through TWMR, 91% showed a directionally concordant effect between the transcript-to-phenotype, transcript-to-metabolite, and metabolite-to-phenotype estimates. Besides validating known and hypothesizing new biological associations, dissection of these causal effects provides clues as to the molecular mechanism through which involved genes modify complex phenotypes. This information is particularly valuable to identify key molecular mediators of highly pleiotropic genetic regions, such as the *TMEM258/FADS1/FADS2* locus (Figure 3). While transcript levels of these genes affected twelve metabolites, two complex lipids were highlighted as strong molecular mediators of the transcript-to-phenotype effects.

Strikingly, 67% of the 206 causal transcript-metabolite-phenotype triplets were missed by TWMR – an approach that only considers gene expression and GWAS data. We highlight two novel but biologically plausible mechanisms linking *ANKH* to calcium levels through modulation of citrate and *SLC6A12* to serum creatinine levels through regulation of the renal osmolyte betaine. Simulation analyses showed that these signals were likely missed by TWMR due to lack of power, as mediation analysis is better suited to detect associations with a low direct to total effect proportion and stronger transcript-to-metabolite than metabolite-to-phenotype effect. Promisingly, our simulations showed that mediation analysis becomes increasingly powerful over a wider range of parameter settings as the sample size of the mediator QTL study increases, highlighting the importance of generating large and publicly available molQTL datasets that can help to unravel functional gene-to-phenotype mechanisms.

As illustrated through the selected examples, a large fraction of detected mediations involves genes encoding metabolic enzymes or transporters/channels, with an enrichment for “secondary active transmembrane transporter activity” (GO:0015291; *FDR* = 0.017 ; background: 7,883 genes with ≥ 3 IVs assessed through TWMR). These results are not surprising given that the proteins encoded by such genes directly interact with metabolites, making it more likely that the effect of changes in their expression are mediated by metabolites. While our method is well-suited to detect such effects, interpretation of discovered mediations is limited by the lack of spatial resolution of the mQTL data. Access to metabolite concentrations in different cellular compartments (e.g., extracellular space, cytosol, mitochondrial matrix, etc.) would generate more fine-tuned mechanistic hypotheses that consider the directionality of metabolite fluxes. Another limitation of our approach is that owing to linkage disequilibrium and regulatory variants affecting multiple genes, transcripts from adjacent genes might appear to be involved in the same signals, as exemplified with the *TMEM258/FADS1/FADS2* locus (Figure 3). While literature supports the role of the *FADS* genes, one cannot exclude a role for *TMEM258*, nor disentangle the specific function of *FADS1* and *FADS2*. Finally, it has been shown that complex phenotypes have a stronger impact on gene expression than the opposite [15]. Due to the lack of *trans*-eQTL data, our method does not investigate reverse causality on metabolites and gene expression, even though accounting for these effects could refine interpretation of the molecular mechanisms shaping complex traits.

In conclusion, we developed a modulable MR framework that has increased power over classical MR approaches to detect causal transcript-to-phenotype relationships when these are mediated by alteration of metabolite levels and is likely to become increasingly powerful upon release of larger molQTL datasets.

## Methods

### Univariable Mendelian Randomization analyses

Transcriptome-wide and metabolome-wide Mendelian randomization (TWMR [12] and MWMR, respectively) were used to estimate the causal effects of transcript and metabolite levels (exposure) on various outcomes. For each transcript/metabolite, using inverse–variance weighted (IVW) method for summary statistics [37], we define the causal effect of the molecular traits on the outcome as

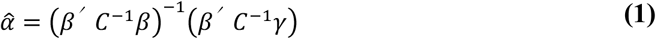

Here, *β* is a vector of length *n* containing the standardized effect size of *n* independent SNPs on the gene/metabolite, derived from eQTL/mQTL studies. *γ* is a vector of length *n* containing the standardized effect size of each SNP on the outcome. *C* is the pairwise LD matrix between the *n* SNPs.

Instrumental variables (IVs) were selected as autosomal, non-strand ambiguous, independent (*r*^*2*^ < 0.01), and significant (*P*_*eQTL*_ < 1.8 × 10^−05^ / *P*_*mQTL*_ < 1.0 × 10^−07^) eQTL/mQTLs available in the UK10K reference panel [38] using PLINK (v1.9) [39]. As retained SNPs are independent, we used the identity matrix to approximate *C*. SNPs with larger effects on the outcome than on the exposure were removed, as these potentially indicate violation of the MR assumptions (i.e., likely reverse causality and/or confounding).

The variance of α can be calculated approximately by the Delta method

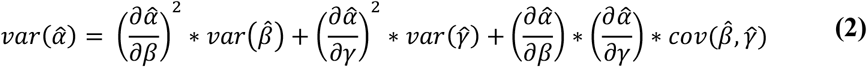

where 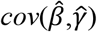 is 0 if *β* and *γ* are estimated from independent samples. We defined the causal effect Z-statistic for gene *i* as 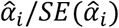, where 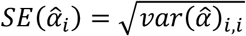.

The IVW method provides an unbiased estimate under the assumption that all genetic variants are valid IVs, i.e., all three MR assumption hold. However, the third assumption (no pleiotropy) is easily violated, leading to inaccurate estimates when horizontal pleiotropy occurs [40]. To test for the presence of pleiotropy, we used Cochran’s Q test [41, 42] to assess whether there were significant differences between the TWMR-derived effect of an instrument on the outcome (i.e., *αβ*_>_) and the GWAS-estimated effect of that instrument on the outcome (*γ*_>_). We defined

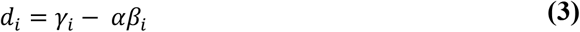

and its variance as

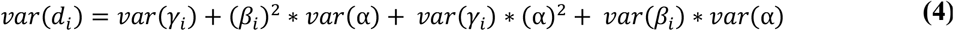

Next, we tested the deviation of each SNP using the test statistic

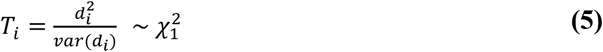

When *P* < 0.05, we removed the SNP with largest |*d*_>_| and then repeated the test.

### Mediation analysis through multivariable Mendelian Randomization analyses

We used a multivariable MR approach to dissect the total causal effect of transcript levels on phenotypes (*α*_*TP*_) into a direct (*α*_*d*_) and indirect (*α*_>_) effects measured through a metabolite. Through inclusion of a metabolite and its associated genetic variants (*r*^*2*^ < 0.01, *P*_mQTL_ < 1 × 10^−07^), the direct effect of gene expression on a phenotype can be estimated using a multivariable regression model [41] as the first element of

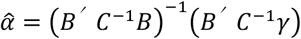

where *B* is a matrix with two columns containing the standardized effect sizes of the IVs on transcript levels in the first column and on the metabolite levels in the second column, *γ* is a vector of length *n* containing the standardized effect size of each SNP on the phenotype, and *C* is the pairwise LD matrix between the *n* SNPs.

### Omics and traits summary statistics

We used eQTL data from the eQTLGen Consortium [3] (N = 31,684), which includes *cis*-eQTLs (< 1 Mb from gene center, 2-cohort filter) for 19,250 transcripts (16,934 with at least one significant cis-eQTL at *FDR* < 0.05 corresponding to *P* < 1.8 × 10^−05^). mQTL data originate from Shin et al. [6], which used ultra-high performance liquid chromatography-tandem mass spectrometry (UPLC-MS/MS) to measure 486 whole blood metabolites in 7,824 European individuals. Association analyses were carried out on ∼2.1 million SNPs and are available for 453 metabolites at the Metabolomics GWAS Server (http://metabolomics.helmholtz-muenchen.de/gwas/). GWAS summary statistics for the 28 outcome traits measured in the UK Biobank (UKB) [43] originate from the Neale Lab (http://www.nealelab.is/uk-biobank/).

### Simulation analyses

We conducted simulation analyses to assess the gain in power upon inclusion of metabolomics data in our MR framework. We simulated a scenario where a transcript has an effect on a phenotype mediated by a metabolite. Two parameters were allowed to vary: the proportion (*ρ*) of direct effect (i.e., effect not mediated by the metabolite) and the ratio (*σ*) between the effect of the transcript on the metabolite (*α*_*TM*_) and of the metabolite on the phenotype (*α*_*MP*_). Other parameters were fixed, including the heritability of the transcript at 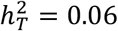 (corresponding to the median *h*^2^ in the eQTLGen data), the number of IVs *N*_*IVs*_ at 6 (corresponding to the median number of IVs used in TWMR analyses). Effect sizes *β*_*eQTL*_ are from a normal distribution 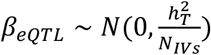. The causal effect of the transcript on the phenotype (*α*_*TP*_) was fixed to 0.035, which results in ∼65% power to detect a significant effect with TWMR. These quantities allowed to define *β*_*GWAS*_ as *β*_*GWAS*_ = *α*_*TP*_ * *β*_*eQTL*_ + *ε*_*P*_, where 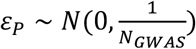 with *N*_*GWAS*_ = 300,000 to reflect the sample size of UKB GWASs.

The same vector of *β*_*eQTL*_ was used to define *β*_*mQTL*_ and estimate the causal effect of the transcript on the metabolite. We defined *β*_*mQTL*_ as *β*_*mQTL*_ = *α*_*TM*_ * *β*_*eQTL*_ *+ ε*_*M*_, where 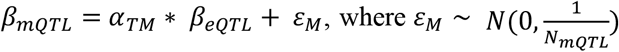 and *N*_*mQTL*_ = 8,000 to reflect the sample size of the mQTL study used in this work and *N*_*mQTL*_ = 90,000 to reflect sample size of potential future studies. We can express the total effect *α*_*TP*_ as *α*_*TP*_ = *α*_*TM*_ * *α*_*MP*_ + *α*_*direct*_, where *α*_*direct*_ represent the direct effect of the transcript on the phenotype and *α*_*TM*_ * *α*_*MP*_ is the indirect effect mediated by the metabolite. Equivalently, *α*_*TM*_ * *α*_*MP*_ = *α*_*TP*_ * (1 − *ρ*) where 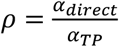. As we are interested in the ratio between the effect of the transcript on the metabolite and the effect of the metabolite on the phenotype (i.e., *σ* = *α*_*TM*_/*α*_*MP*_), we can express *α*_*TM*_ as 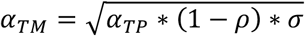. Similarly, to estimate the effect of the metabolite on the phenotype, we considered a metabolite with heritability 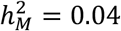 (corresponding to the median of *h*^2^ in the KORA+TwinsUK mQTL data) and *N*_*IVs*_ = 5 (corresponding to the median number of IVs used in MWMR analyses). Effect size *β*_*mQTL*_ are from a normal distribution 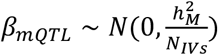. These quantities allowed to define *β*_*GWAS*_ as 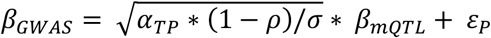, where 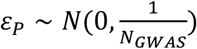. Ranging *ρ* and *σ* fro 0 to 1 and from 0.1 and 10, respectively, we run each simulation 1,000 times and performed TWMR and MWMR starting from above-described *β*_*eQTL*_, *β*_*mQTL*_ and *β*_*GWAS*_. For each MR analysis we calculated the power to detect a significant association and plotted the difference in power between TWMR and the mediation analyses (i.e., *power*_*TP*_ − *power*_*TM*_ * *power*_*MP*_) as a heatmap.

## Data and code availability

All data used in this study is publicly available (see Web resources). Produced data is available as supplemental tables. Code used to perform analyses is freely available at https://github.com/eleporcu/Gene_Metab_Pheno.

## Author contributions

E.P. and Z.K. conceived and designed the study; E.P., M.C.S. and Z.K. contributed to the mathematical derivations of the research; E.P. performed statistical analyses; C.A. and A.R. contributed with the biological interpretation of the results; E.P., C.A. and Z.K. drafted the manuscript; M.C.S. and A.R. revised the manuscript. All authors read the paper and contributed to its final form.

## Acknowledgments

Computations were carried out on the high-performance cluster of the Lausanne University Hospital (CHUV). This work was supported by funding from the Department of Computational Biology (Z.K.) and the Center for Integrative Genomics (A.R.) from the University of Lausanne, as well as funding from the Swiss National Science Foundation (310030-189147 to Z.K. and 31003A_182632 to AR).

## Web resources

- eQTLGen data https://www.eqtlgen.org
- GWAS Catalog https://www.ebi.ac.uk/gwas/
- GWAS UK Biobank summary statistics from Neale’s Lab http://www.nealelab.is/uk-biobank/
- Metabolomics GWAS Server http://metabolomics.helmholtz-muenchen.de/gwas/
- UCSC Genome Browser https://genome.ucsc.edu
- STRING database https://string-db.org/

