## Supplemental Figures for "Exploiting the mediating role of the metabolome to unravel transcript-to-phenotype associations"

**Supplemental Figure 1.** Distribution of σ (i.e. the ratio between the transcript-to-metabolite and the metabolite-to-phenotype effects) for the 206 casual triplets.

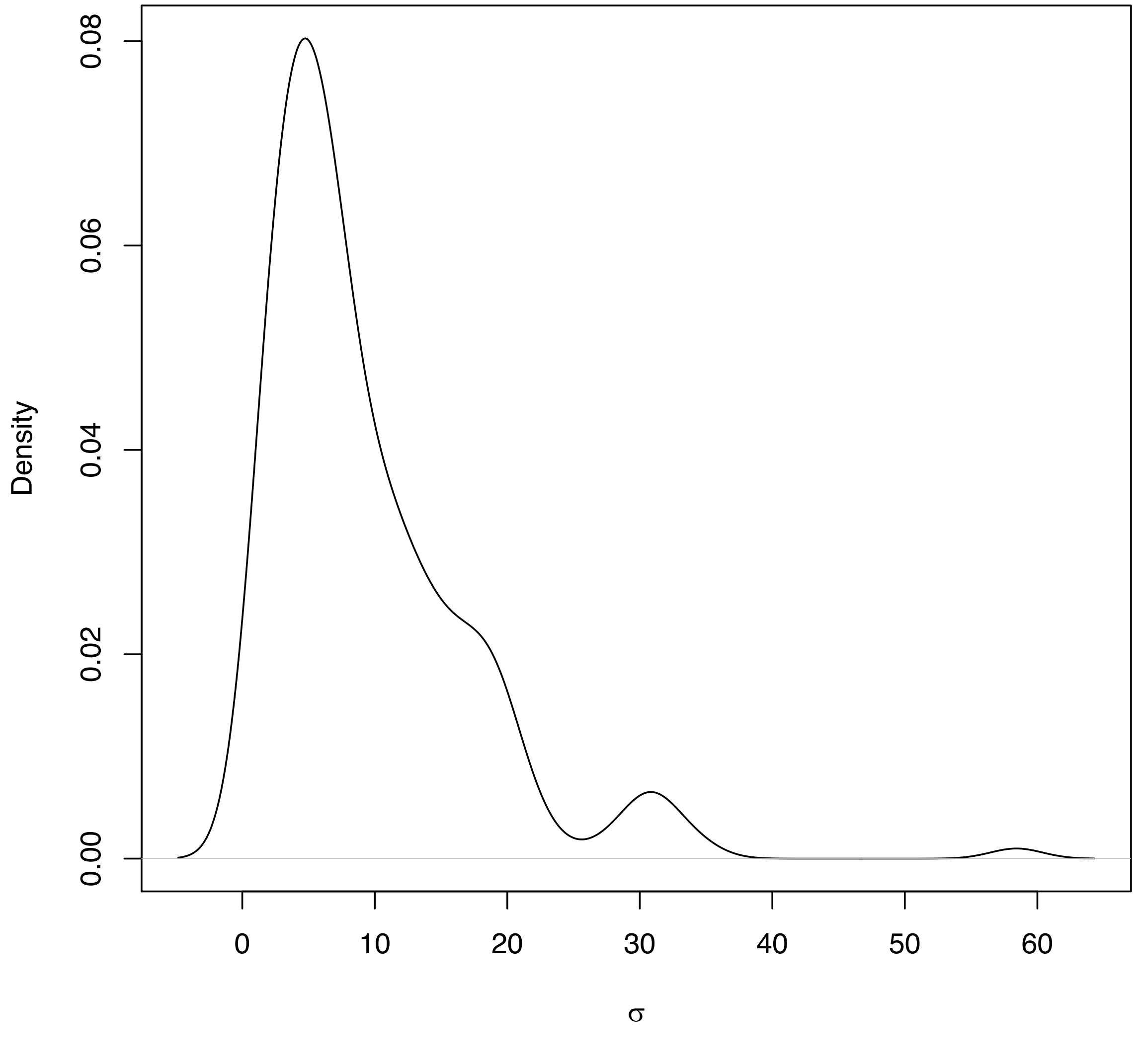
